# Spatial Analysis of Malignant-Immune Cell Interactions in the Tumor Microenvironment Using Topological Data Analysis

**DOI:** 10.1101/2025.03.27.645762

**Authors:** Seol Ah Park, Yongsoo Kim, Paweł Dłotko, Davide Gurnari, Jooyoung Hahn

## Abstract

The spatial interactions between malignant and immune cells in the tumor microenvironment (TME) play a crucial role in cancer biology and treatment response. Understanding these interactions is critical for predicting prognosis and assessing immunotherapy effectiveness. Conventional methods, which focus on local spatial features, often struggle to achieve robust analysis due to the complex and heterogeneous cellular distributions. We propose a Topological Data Analysis (TDA)-based framework using both global and local spatial features between malignant and immune cells. For the global aspects, we introduce Topological Malignant Clusters (TopMC), a method utilizing persistence diagrams in TDA to capture the global spatial shapes of malignant cell distributions. It quantifies tumor-immune cell infiltration at a global scale by computing distances from the boundaries of the TopMC. Local interactions are evaluated through the density of malignant cells. Using high-resolution multiplex immunofluorescence (mIF) imaging in Diffuse Large B Cell Lymphoma, we integrate distance and density measures into a distance-density space to analyze patterns of malignant-immune cell interactions in TME. This study shows the robustness of the proposed approach to variations in cell distribution, enabling consistent analysis irrespective of whether images are acquired from malignant-enriched or border regions of the tumors. Furthermore, we elucidate the correlation between spatial patterns of immune phenotypes and patient survival probability.

## 1 Introduction

Elucidating the spatial interactions between malignant and immune cells in the tumor microenvironment (TME) is critical for understanding tumor progression, predicting prognosis, and the patient outcomes [Arneth, 2019, Binnewies et al., 2018, Sadeghi Rad et al., 2021]. The spatial distribution of these cells within the TME reflects complex biological processes including immune infiltration and evasion. Spatial analysis of tumor and immune cells in the TME may enable us to quantitatively disentangle the underlying biological processes, which we can link to patient outcome [Chen et al., 2024, Elhanani et al., 2023, Fu et al., 2021].

Numerous studies have developed methods to extract spatial features from cellular imaging data to correlate these features with patient outcomes. For instance, spatial features related to immune cell infiltration in breast cancer have been quantified using point cloud data and rasterized grid images, including cell counts, densities, and statistical measures with nearest-neighbors such as the *G* and cross-*G* functions [Page et al., 2023]. A graph-based method has represented cellular neighborhoods as graphs to extract meaningful features that enable patient clustering and comparative analyses [Ehsani et al., 2023]. Another framework has outlined five essential spatial analysis aspects: localization, structure-based, distance-based, graph-based communication, and neighborhood analyses, highlighting their potential as clinical diagnostic metrics [Semba and Ishimoto, 2024]. Similarly, spatial and non-spatial features have been combined in diffuse large B-cell lymphoma, demonstrating complementary prognostic insights from methods such as Ripley’s *L* function alongside traditional cell counts and densities [Roemer et al., 2023].

Despite these advances, existing approaches predominantly focus on local cell-to-cell interactions (e.g., shortest cell distances, nearest-neighbor measures, Ripley’s functions). Although graph-based or rasterized grid methods partially reduce locality constraints, they still emphasize local metrics through node connectivity or predefined regions. Such local-centric methods often struggle to provide robust analyses for highly heterogeneous or structurally complex tumors.

To address these limitations, we propose a framework based on Topological Data Analysis (TDA), integrating both global and local spatial features for a comprehensive understanding of interactions between malignant and immune cells. TDA robustly handles high-dimensional data with inherent noise [Amézquita et al., 2020, Loughrey et al., 2021, Skaf and Laubenbacher, 2022], effectively summarizing complex information and thus outperforming traditional statistical methods. TDA has uncovered clinically meaningful disease subgroups in genomic studies [Li et al., 2015, Nicolau et al., 2011]. Beyond genomics, TDA has also shown effectiveness in image segmentation across various imaging modalities [Belchi et al., 2018, Lukasczyk et al., 2020, Pike et al., 2020]. Furthermore, persistent homology, a component of TDA, enables quantitative assessment of tissue morphology across different cancer stages [Lawson et al., 2019, Singh et al., 2014], offering insights into cancer progression.

Among TDA tools, we utilizes **Ball Mapper** [Dłotko et al., 2024], a variant of the Mapper algorithm [Singh et al., 2007], to effectively visualize high-dimensional data and capture continuous changes in spatial features. Compared to the classical Mapper, Ball Mapper requires fewer parameters, specifically only a distance metric and ball radius, and takes less computational costs.

In this study, we introduce Topological Malignant Clusters (TopMC), designed to capture global spatial patterns of malignant cell distributions. Unlike traditional methods that focus on cell-to-cell distances, TopMC utilizes cell-to-cluster distances, enhancing robustness against measurement noise. Moreover, TopMC effectively characterizes variations in cellular configurations, enabling comprehensive analysis of complex and heterogeneous spatial patterns in the TME. Based on the computed TopMC, we propose a framework for understanding malignant-immune cell interactions, focusing on immune cell infiltration and the density of malignant cells with diverse configurations. In order to examine these interactions, we use point cloud data of high-resolution multiplex immunofluorescence (mIF) imaging in diffuse large B cell lymphoma (DLBCL). The images are annotated into two types: the malignant-enriched or border regions of the tumors. Regardless of two types of images, TopMC can capture various global spatial shapes of malignant cell distributions. It eventually brings further analysis to reveal spatial characteristics of two types of images and hidden patterns of malignant-immune cell interactions linked to clinical outcomes.

## 2 Materials and Methods

### 2.1 Data Acquisition

In this study, we use the patient data in [Roemer et al., 2023] and point clouds in [Kim et al., 2023] from mIF imaging of primary central nervous system lymphoma patients. The images are annotated independently into two types by pathologists on a sample-by-sample basis ensuring no patient-specific bias:

- TR image: the malignant-enriched regions of the tumors.
- BR image: the border regions of the tumors.

These regions are selected by pathologists based on qualitative criteria, such as malignant cell infiltration; TR images show high malignant cell concentrations, while BR images illustrate interactions with adjacent cerebral tissue.

From given point clouds in the domain Ω of mIF imaging data, let us denote locations of malignant cells (PAX5+) as reference point cloud ℛ and locations of immune cells of the *i*^th^ aggregated phenotype as query point cloud 𝒬^*i*^:

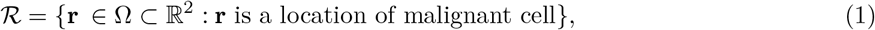

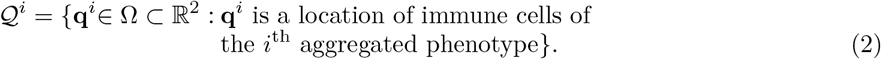

where

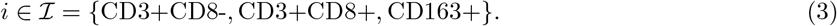

In the following subsections, we describe the method of constructing the Topological Malignant Clusters (TopMC) from the reference point cloud ℛ. We then detail the extraction of two key features:

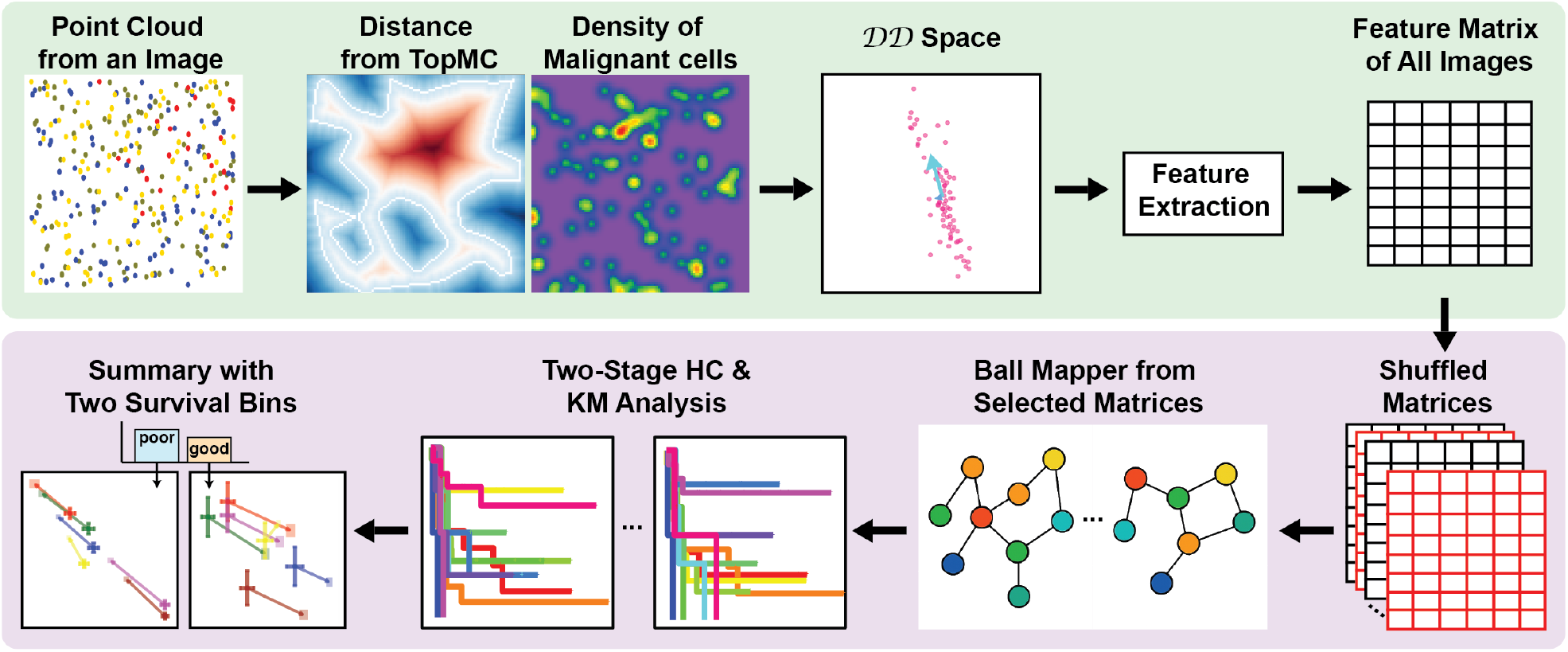

**Graphical Abstract of the Proposed Framework** For each point cloud, we compute distance functions from the boundaries of Topological Malignant Clusters (TopMC) and density functions from malignant cell distributions. These paired values at immune cells form the distance-density (𝒟 𝒟) space, from which spatial features are extracted to build a comprehensive feature matrix. To explore diverse perspectives of the high-dimensional features, we repeatedly shuffle the matrix, selecting statistically significant shuffled matrices via optimal cluster identification. From graphs of Ball Mapper, we further analyze through a two-stage hierarchical clustering (HC), followed by Kaplan-Meier (KM) survival analysis on identified patient clusters. All valid patient clusters are collected by corresponding survival probabilities and spatial features from selected matrices. Survival probabilities are divided into two survival bins, with spatial features averaged within each bin to identify associations between spatial interaction patterns and clinical outcomes.

- Quantification of the infiltration of immune cells 𝒬 ^*i*^ into the TopMC.
- Quantification of the local concentration of malignant cells ℛ around immume cells 𝒬 ^*i*^.

### 2.2 Topological Malignant Clusters (TopMC)

Given reference points ℛ in (1), we use TDA with the Vietoris-Rips complex and persistence diagrams (PD) [Carlsson, 2009, Edelsbrunner and Harer, 2010, Ghrist, 2008, Zomorodian, 2005] to capture global spatial shape of malignant cells, illustrated in Fig. 1-(c), where the olive-colored regions (TopMC) mostly overlap with the regions of malignant cells (coral red) in Fig. 1-(a). The PD summarizes topological features (connected components or loops) over varying scale lengths. Ideally, the PD should cover all scale lengths, but this becomes computationally challenging due to the large number of points in each image and the extensive quantity of images processed [Glisse et al., 2014–2024]. Hence, we set a practical limit on the maximum scale length:

**Figure 1:**
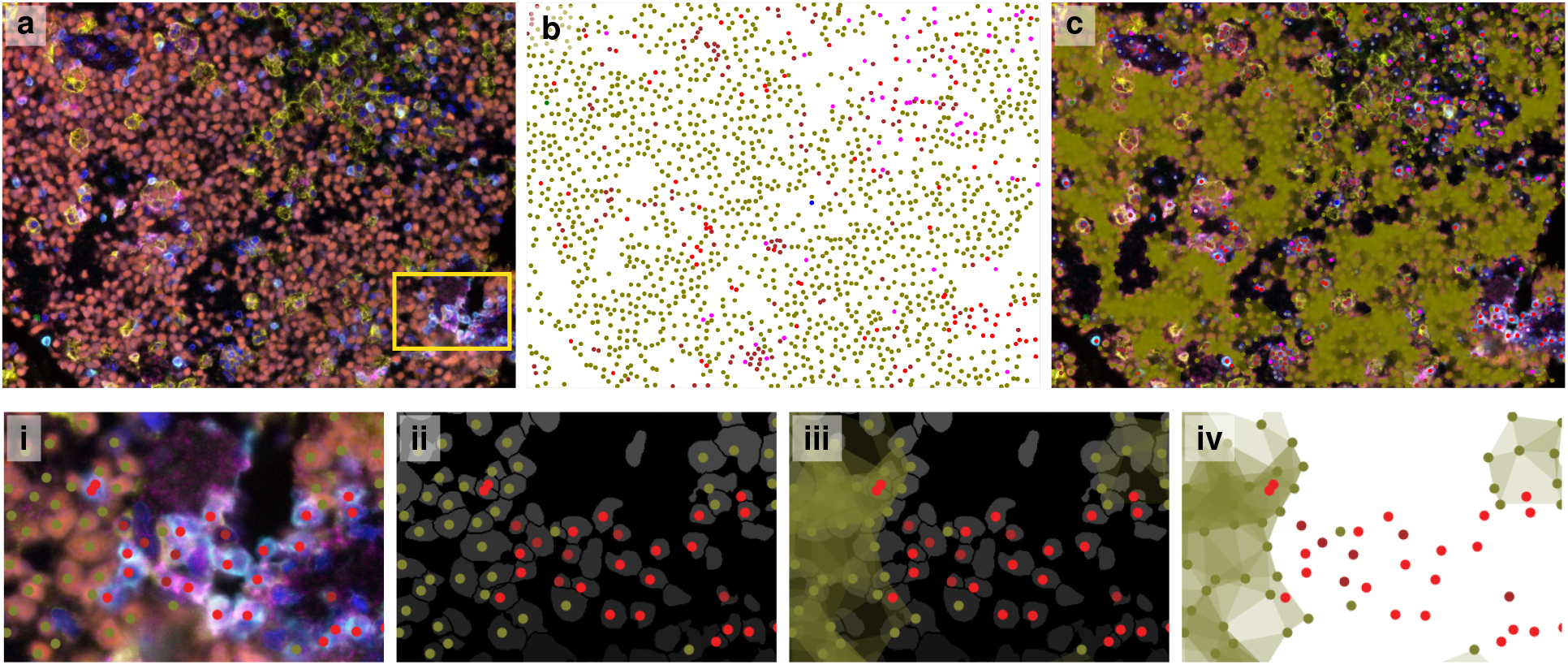
TopMC: Global Spatial Shapes of Malignant Cell Distributions: **(a)** A raw multiplex immunofluorescence (mIF) image as shown in [Roemer et al., 2023]. The highlighted area by the yellow rectangle is presented with more details in the second row. **(b)** Point clouds representing cell centers, color-coded by phenotype as listed in Table 1. **(c)** TopMC presented by olive-colored area, overlaid on (a), qualitatively demonstrates glboal spatial shapes of malignant cell distributions in (a) and (b). **(i)** illustrates different phenotypes on highlighted area in (a). **(ii)** shows the segmented image with cell centers. **(iii)** presents the TopMC on (ii). **(iv)** illustrates the spatial context between immune cells and TopMC, delineating an interaction.

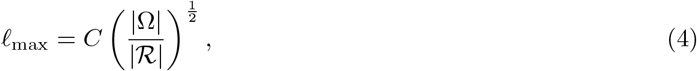

**Table 1:** Color Legend: Summary of cell types, phenotypes, aggregated phenotypes, and their associated colors.

| Cell Type | Phenotype | Color | Aggregated Phenotype | Color |
| --- | --- | --- | --- | --- |
| T cells (CD3+) | Naive (CD8-), PD1-<br>Naive (CD8-), PD1+ | red<br>blue | CD3+CD8- | indigo |
|  | Active (CD8+), PD1-<br>Active (CD8+), PD1+ | green<br>yellow | CD3+CD8+ | deep pink |
| Malignant cell | PAX5+ | olive | PAX5+ | olive |
| Macrophage (CD163+) | PDL1-<br>PDL1+ | magenta<br>brown | CD163+ | gold |

where |ℛ| is the number of points in ℛ, |Ω| is the area of the mIF image domain Ω, and *C* = 3. It defines a scale that represents an average distance between points in a 2D space. The constant *C* is a choice for balancing computational efficiency with the need to capture more topological features.

Now, we obtain the PD using *ℓ*_max_ in (4). Each point (*b, d*) in PD marks the birth and death of a topological feature. We select the most persistent feature (the dominant topological structure in ℛ) by the longest finite lifetime in the PD:

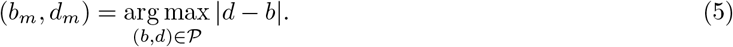

To define a biologically plausible characteristic length, we take the midpoint between *b*_*m*_ and *d*_*m*_ and constrain it by the biologically relevant separation 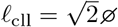, where ∅ ≈ 50*μm* is the average diameter of a malignant cell [Hao et al., 2018]:

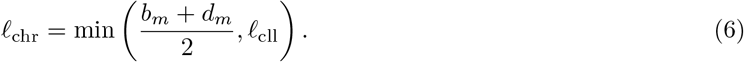

Using this characteristic length, we reconstruct the global spatial shape of malignant cells by the union of 2D simplices (triangles), which we refer to as **topological malignant clusters (TopMC)**.

### 2.3 Feature Matrix

In the following subsections, we descirbe a process of obtaining the feature matirx and its graphical overview is illustrated in Fig. 2 or 4.

**Figure 2:**
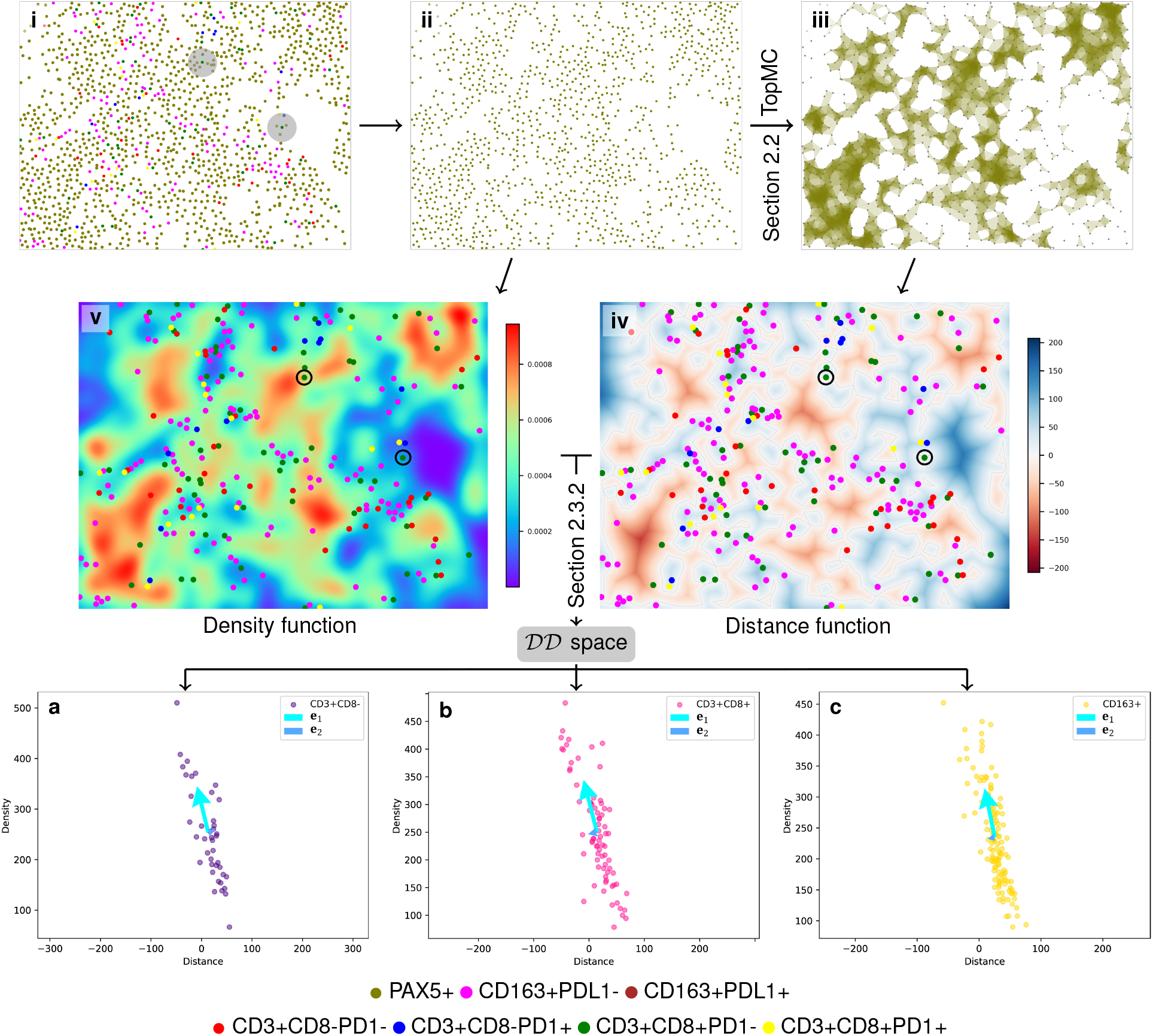
Distance and Density Functions of Point Clouds from the Malignant-Enriched Regions of Tumors (TR image): **(i)** shows the original point cloud in TR image. **(ii)** presents only malignant cell distributions. **(iii)** shows TopMC from (ii). **(iv)** is the signed distance function from the boundaries of TopMC with immune cells overlaid, illustrating the varying degrees of immune cell infiltration. **(v)** is the density function that evaluates the local concentration of malignant cells around the immune cells. Regarding two immune cells marked by gray areas in (i) and black circles with dotted lines in (iv) and (v), see detailed explanation in Section 3.2. **(a), (b)**, and **(c)** show the 𝒟𝒟^*i*^ spaces in (10) for *i* ∈ ℐ in (3). We also demonstrate 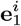 and 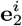at **m**^*i*^ from (11) in the 𝒟𝒟^*i*^ space, considered as a visualization of 18D feature vector in (12).

**Figure 3:**
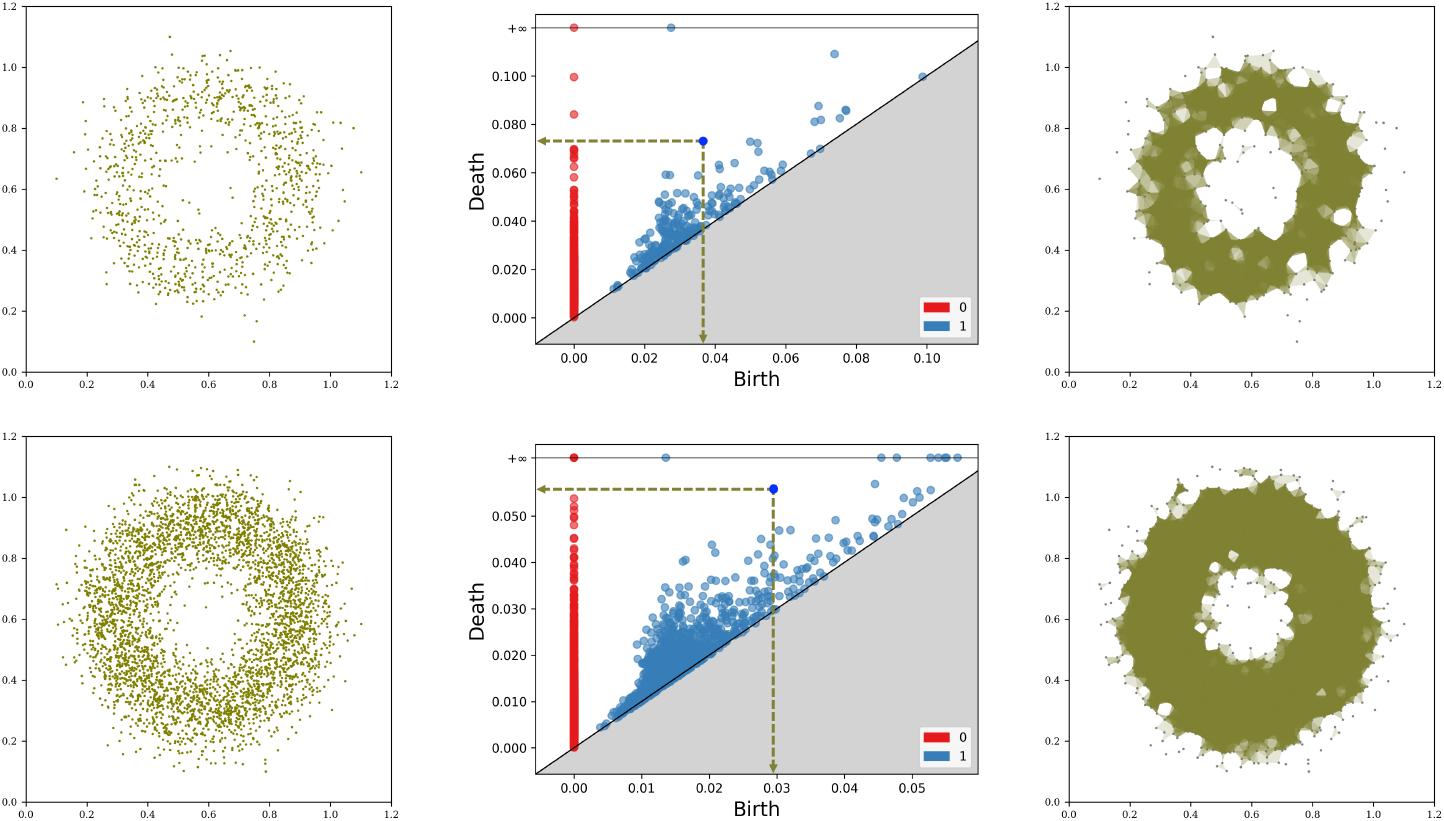
Persistence Diagrams and TopMC for Point Clouds with Large Density Difference: The left column shows point clouds with 1 10^3^ (top) and 4 10^3^ (bottom) points in the same domain. The middle column presents the corresponding persistence diagrams, computed by the maximum scale length in (4), *ℓ*_max_ = 1.14 · 10^*−*1^ (top) and 5.69 · 10^*−*2^ (bottom). From the most persistent features in the PDs in (5), (3.66 10^*−*2^, 7.30 · 10^*−*2^) (top) and (2.95 · 10^*−*2^, 5.58 · 10^*−*2^) (bottom), the characteristic length *ℓ*_chr_ in (4) is calculated with with *ℓ*_cll_ = ∞ in these examples. The right column visualizes the corresponding TopMCs, which robustly show an annulus shape.

#### 2.3.1 Distance Function from TopMC

Let us denote 𝒯 as TopMC of ℛ in (1). Then, the solution of the eikonal equation provides a distance function from *∂*𝒯, the boundary of 𝒯 :

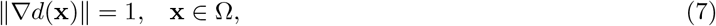

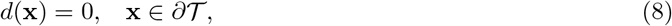

where ∥·∥ is the Euclidean norm in ℝ^2^, computed by the fast marching method [Sethian, 1999] and scikit-fmm Python library [Furtney]. In order to quantify the infiltration of immune cells 𝒬^*i*^ into the TopMC, we measure a signed distance from *∂*𝒯 :

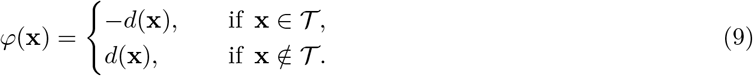

See the signed distance function *φ* from TopMC in Fig. 2 or 4(iv).

#### 2.3.2 Distance-Density Space

We quantify local concentrations of malignant cells around immune cells using a Gaussian-based density function. For reference points in ℛ in (1), we calculate density within a local region and combine densities across all points using Gaussian kernel convolution. To analyze density jointly with spatial relationships (signed distance in (9)), we rescale the density to match the signed distance range based on global extremes; See the density function *Ψ* (A1.3) of malignant cells in Fig. 2 or 4(v). Detailed formulation is provided in Appendix A1.

Now, we define the distance-density 𝒟𝒟 space, for *i* ∈ ℐ in (3):

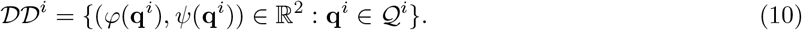

The points in the ^*i*^ space are illustrated in Fig. 2 and 4(a), (b), and (c). The measures, distance *φ*(**q**^*i*^) and density *Ψ*(**q**^*i*^), are complementary yet independent, capturing distinct aspects of malignant-immune interactions. While the cell-to-cluster distance indicates immune cell positions relative to the boundary of TopMC, the density measure reflects the local concentration of malignant cells around immune cells.

**Figure 4:**
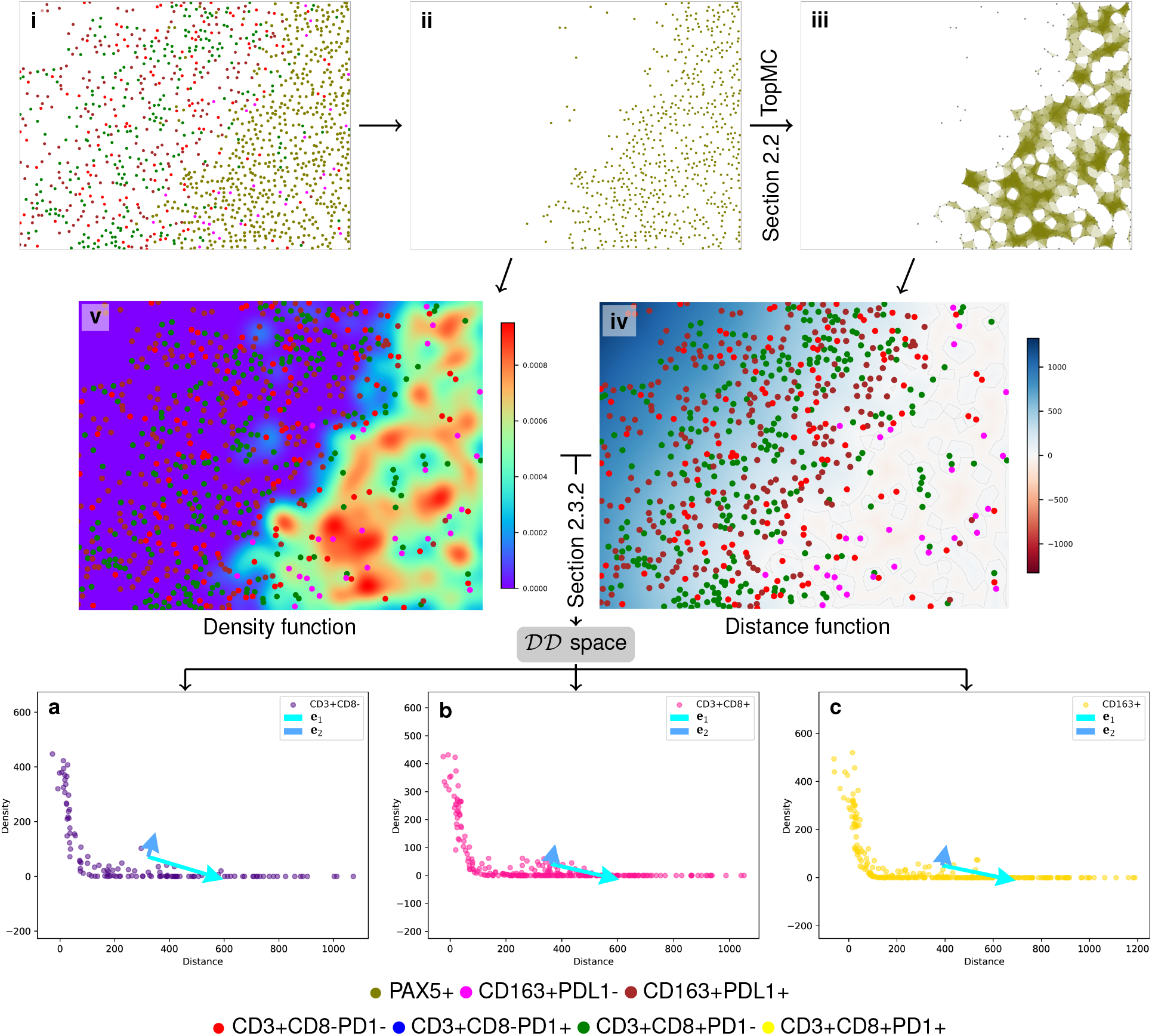
Distance and Density Functions of Point Clouds from the Border Regions of Tumors (BR image): The scenarios of Fig. 2-(i) from TR image and (i) from BR image above are standard examples of strong and weak interaction with malignant cells, respectively; see more details in Section 3.2. All procedures in figures are same as Fig. 2.

In order to effectively summarize three distance-density spaces, we use the principle component analysis (PCA) on 𝒟𝒟^*i*^ and define a row vector:

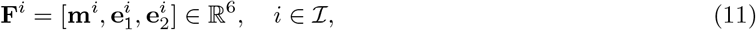

where 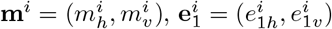, and 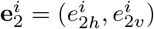 are 2D vectors to indicate averages of distance and density values in 𝒟𝒟^*i*^ space, the first and the second principle components of 𝒟𝒟^*i*^, respectively. Finally, for the *n*^th^ image, we define the row vector of the feature matrix by the concatenation of three row vectors:

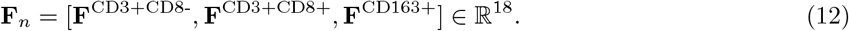

Note that the row vector from a given mIF image is only available if |*Q*^*i*^| ≥ 2 for all *i*∈ ℐ. Based on this criterion, 972 mIF images out of the 1337 images in the DLBCL study are analyzed, including 687 TR images and 285 BR images.

### 2.4 Graph Representation of High-Dimensional Data

We employ Ball Mapper (pyBallMapper, [Gurnari]) to visualize high-dimensional data from mIF images, 18-dimensional (18D) feature in (12) captures the infiltration of immune cells and the local concentration of malignant cells around immune cells. Using the *cosine dissimilarity* and the parameter *ϵ* of the spatial granularity, Ball Mapper represents this data as a simplified graph with nodes and edges. A node consists of row vectors in the feature matrix in (12) with similar characteristics based on the cosine dissimilarity and an edge between nodes indicates that there is at least one common feature shared by nodes. Adjusting *ϵ* allows control over the resolution of the graph, with smaller values producing a more detailed graph and larger values generating broader connections among features. In Section 3.3, we show the the graph of Ball Mapper efficiently identifies patterns and relationships in 18D features, as explored in our study with *ϵ* = 4.5 · 10^−2^.

### 2.5 Optimal Cluster Identification via Two-Stage HCs

We use the graph of Ball Mapper to quantify immune cell activity and its relationship with survival outcomes in DLBCL patients. Given the varying number of images per patient (from 3 to 48), we adopt a two-stage hierarchical clustering (HC) approach. Using the feature matrix in (12), Ball Mapper visualizes the 18D space features as a graph in Fig. 5. While effective, it does not capture every aspect of the high-dimensional data. To explore the data from multiple perspectives, we randomize the row order in the feature matrix and reapply Ball Mapper multiple times. This allows us to examine the data from multiple angles without altering the data itself in high-dimensional space. Through multiple observations, we correlate the observed patterns in the images with survival rates in DLBCL patients, identifying statistically significant variations.

**Figure 5:**
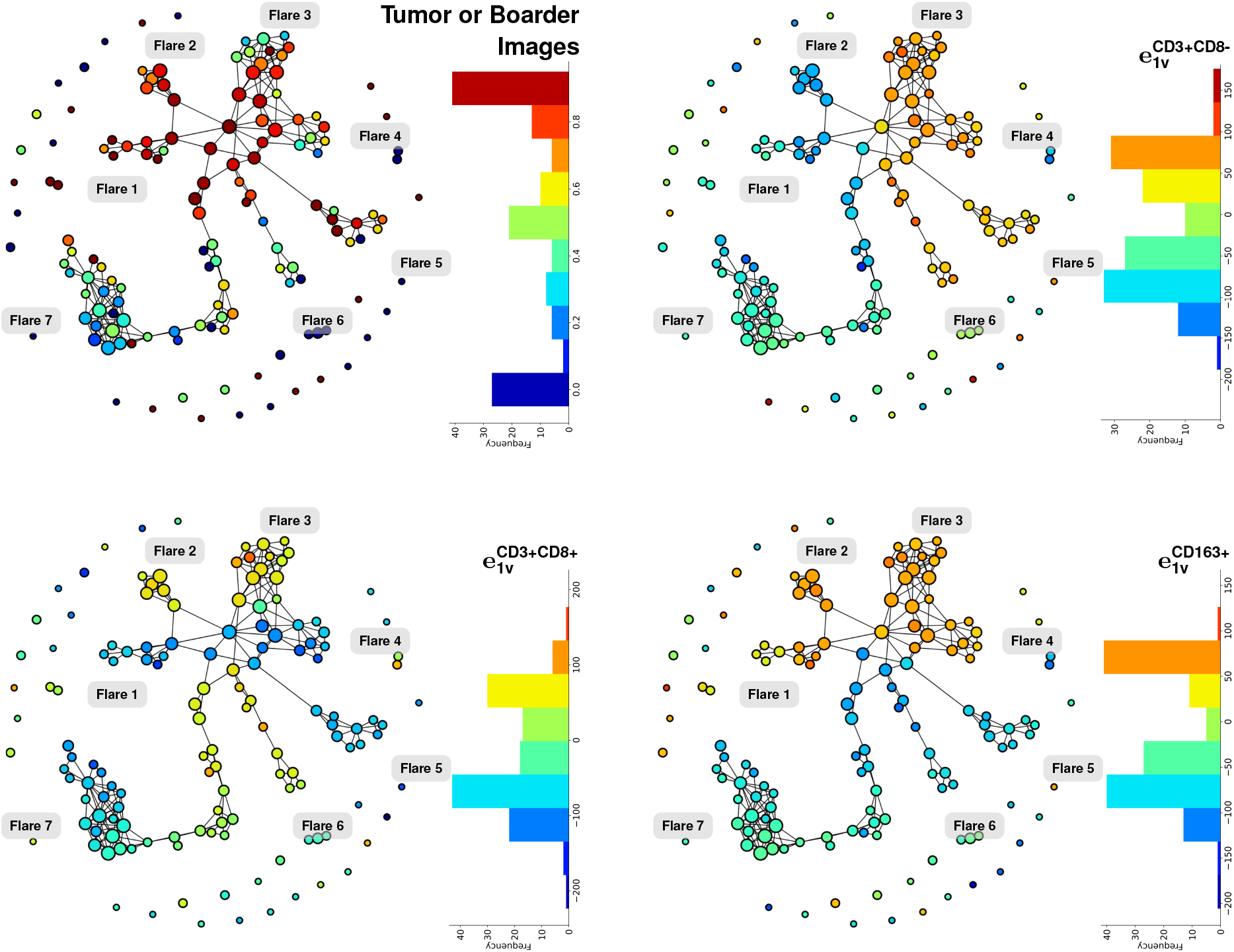
Ball Mapper of Feature Matrix: The graphs of Ball Mapper illustrate assigned scalar values on images. In the top-left panel, TR and BR images are assigned values of 1 and 0, respectively, and each node is colored by the average of these values. As a result, red (jet scale) represents nodes of TR images, while blue corresponds to BR images. In the 18D space, this graph summarizes how TR and BR images are distributed based on similar features in (12). Using the same coloring method, the scalar attribute of 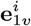for *i* ∈ ℐ is visualized in the remaining panels; see more details in Section 3.3. Next to each graph, the distribution of the corresponding value is presented.

To perform this analysis, we generate |*J*| = 1000 feature matrices by shuffling rows of the feature matrix based on uniform distribution, where *J* is an index set of shuffled matrices. Given a pair (*K, L*) of positive integers, we repeat the following procedure for each *j*^th^ matrix, *j* ∈ *J*. A graph is computed by Ball Mapper with the same parameter *ϵ*. On each node in the graph with *N*_*j*_ nodes, averaging feature vectors ((12)) in the node, we construct a 18 × *N*_*j*_ matrix. We then apply the first stage of HC with *K* clusters to the obtained matrix. In the second stage, we construct a *K* × *Q* matrix, where *Q* is the number of patients. Each column vector in this matrix is obtained by counting and normalizing the number of images per patient assigned to each of *K* clusters. We then apply HC with *L* clusters to the *K* × *Q* matrix and compute survival probabilities with event-free survival rates across the *L* clusters using the Kaplan-Meier (KM) analysis. To achieve a statistically meaningful analysis, we count the number of matrices *j* ∈ *J* whose KM analysis yields a *p*-value *p*_*j*_(*K, L*) ≤ *δ*(= 0.05):

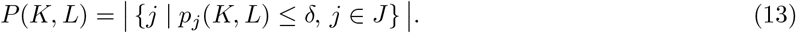

To ensure reliable interpretation of survival-related features, we consider only patient clusters containing at least 10 patients, referring to these as *valid clusters*. For each matrix *j* ∈ *J*, we count the total number of patients belonging to these valid clusters, denoted by *s*_*j*_(*K, L*). We then sum these patient counts across all matrices, *S*(*K, L*) = Σ_*j*∈*J*_ *s*_*j*_(*K, L*); see the algorithm in Appendix A2.

Finally, we determine the optimal numbers of clusters *K*^∗^ and *L*^∗^ by maximizing the statistical significance and robustness of patient grouping as follows:

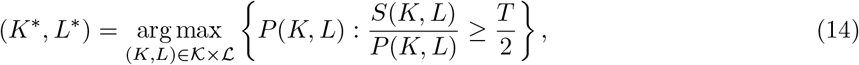

where 𝒦= {5, 6, …, 25},ℒ = {3, 4, …, 25}, *T* is the total number of patients. In two-stage HCs, (*K*^∗^, *L*^∗^) finds the clustering arrangement that makes the analysis statistically robust and clinically meaningful for at least half of the patient population.

## 3 Results

### 3.1 Global Spatial Shapes of Malignant Cells

TopMC in Section 2.2 captures the global spatial shapes of malignant cell distributions from mIF images. It is robust to errors in cell segmentation and consistent regardless of cell densities because of designed characteristic length in (6); see Fig. 3. We use this global spatial shape to explore the intricate malignant-immune cell interactions. In Fig. 1, an mIF image from the DLBCL study presents an example of TopMCs. The TopMC is predominantly occupied by malignant cells, showing a clear boundary between tumor-enriched area and empty space. In a highlighted area by the yellow rectangle in Fig. 1-(a), while most immune cells (CD3+CD8-PD1- and CD163+PDL1+) are located outside TopMC, the panel (iv) shows some immune cells penetrating into TopMC, indicating some degree of infiltration.

### 3.2 Interaction Between Immune and Malignant Cells

In Fig. 2 and 4, we present the signed distance and density functions in (10) for two example images from DLBCL study: tumor-enriched regions (TR image) and border regions (BR image) of the tumors, which have large density difference of malignant cells and heterogeneous configurations of immune cells. The proposed method summarizes such spatial variations of DLBCL images into the 18D space.

The signed distance functions in Fig. 2-(iv) and 4-(iv) show the spatial positioning of immune cells relative to the TopMC boundary, with red indicating negative values (deeper infiltration) and blue showing positive values (greater distance). White areas denote proximity to the boundary. This is complemented by the local density of malignant cells around immune cells, represented in a purple-to-scarlet color bar where scarlet signifies high density, indicating active malignant regions. These two measures, density and signed distance, are used to quantify the extent of immune cell infiltration and their interaction with malignant cells. While they generally correlate, discrepancies can occur, such as immune cells at similar infiltration levels exhibiting varying densities; see the two points marked by small gray disks in Fig. 2-(i). This is illustrated by two active T cells (PD1-) marked by black circles in Fig. 2-(iv), where two immune cells at the boundary show markedly different densities in Fig. 2-(v), underscoring how these measures provide a nuanced view of immune-malignant cell interactions.

For two distinct configurations of malignant cells, we present the results of (10) in the 𝒟𝒟 space (bottom row of Fig. 2 for TR image and Fig. 4 for BR image). In the TR image, we observe a strong interaction between immune and malignant cells, reflected as a nearly linear pattern in the 𝒟𝒟 space, typical for TR images. Conversely, in the BR image, immune cells appear enriched at larger distances from the TopMC with sparse malignant cells nearby, corresponding to an exponential-decay pattern in the 𝒟𝒟 space. Both patterns are effectively described by 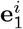 in (11): vectors pointing toward the top-left indicate strong interactions, typical of TR images, while vectors pointing toward the bottom-right imply weak interactions, typical of BR images. However, this interpretation does not always hold because cell distribution can change continuously to diverse and atypical configurations. To better capture the continuous and complex transition of immune-malignant cell interactions, we employ Ball Mapper approach in the next section.

### 3.3 Ball Mapper-Based Summarization of Spatial Patterns

The features in (12) summarize interactions between malignant and all aggregated immune cells in high-dimensional space. We employ Ball Mapper with cosine dissimilarity to visualize these interactions across multiple immune cell types without reducing feature dimensions. In Fig. 5, a node in the graph of Ball Mapper represents a group of images with similar patterns in the 𝒟𝒟 space, quantified by the cosine dissimilarity and the parameter *ϵ*. Edges indicate shared features between nodes. In our analysis of DLBCL images, we primarily focus on the largest connected component in the graph to identify dominant interaction patterns for subsequent analysis.

We elaborate more details about the meaning of the graphs in Fig. 5. For convenience, we refer to the seven visually distinct branches deviating from the main collection of nodes as *flares*. The color on a node means the average of assigned scalar values on rows (images) in the node. For example, on the top-left panel, we assign the scalar value as 0 or 1 for BR and TR images, respectively. The red node contains all TR images, while the blue node contains all BR images. The nodes of the colors between red and blue in the jet color scale mean a mixture of TR and BR images. Now, it is clear to see that Flares from 1 and 5 mainly consist of TR images, though not exclusively.

In three other panels of Fig. 5 excluding the top-left panel, we assign the scalar value as 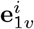 for *i* ∈ ℐ and then let us focus on nodes in the Flare 3 from all panels, simultaneously. Examining the red nodes in Flare 3 (top-left panel), we find these nodes predominantly classified as TR images. Correspondingly, their associated panels indicate large magnitudes of 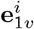, suggesting a nearly linear pattern in the 𝒟𝒟 space. However, we observe that some nodes exhibiting similar patterns also include BR images. It demonstrates that both BR and TR images can share comparable patterns under the cosine dissimilarity measure. In the case of Flare 2, the nodes consist of mostly from TR images, but 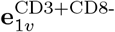 show very different value than the other two immune cells. In other words, even though two images from Flares 2 and 3 are annotated by TR image, the spatial patterns of immune cells in the 𝒟𝒟 space can be different. These observations verify that Ball Mapper provides continuous transitions of malignant-immune cell interactions in DLBCL patient, considering simultaneously multiple immune cell types.

### 3.4 Survival-Linked Spatial Patterns of Immune Cell Activity

We analyze patient clusters derived in Section 2.5, incorporating both TR and BR images. By associating valid patient clusters (clustered by *L*^∗^) with their dominant node clusters (clustered by *K*^∗^), we examine patient clusters in terms of their extracted features within the 𝒟𝒟 space and correlate these with survival probabilities. Here, *T* = 78 in (14) with optimal number of clusters, (*K*^∗^, *L*^∗^) = (8, 12). For illustrative purposes, HC results from a matrix meeting the condition *p*_*j*_(*K*^∗^, *L*^∗^) ≤ *δ* are shown in Fig. 6. The first HC identifies 8 Node Clusters (NCs), differentiated by averaged 18D features in Fig. 6A. The second HC further distinguishes these into 12 Patient Clusters (PCs), as depicted in Fig. 6B, with corresponding survival probabilities shown in Fig. 6C. Among these PCs, *PC*_1_ and *PC*_9_, each containing more than 10 patients, are selected for the analysis. Their node distributions in Fig. 6B show *PC*_1_ predominantly linked to *NC*_1_ and *PC*_9_ to *NC*_7_. We compute averaged 18D features for these dominant node clusters, associating them directly with survival probabilities at the latest time in the KM plot. By tracing the images corresponding to each patient cluster, we calculate averaged values of **m**^*i*^ and 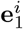 for each *i* ∈ ℐ. For all selected feature matrices satisfying *p*_*j*_(*K*^∗^, *L*^∗^) ≤ *δ*, we collect associated survival probabilities along with averaged feature values across phenotypes. Finally, we summarize the relationship between immune cell activity and survival probability by dividing survival data into two bins and visualizing these findings in the *mean* 𝒟𝒟 space (Fig. 7).

**Figure 6:**
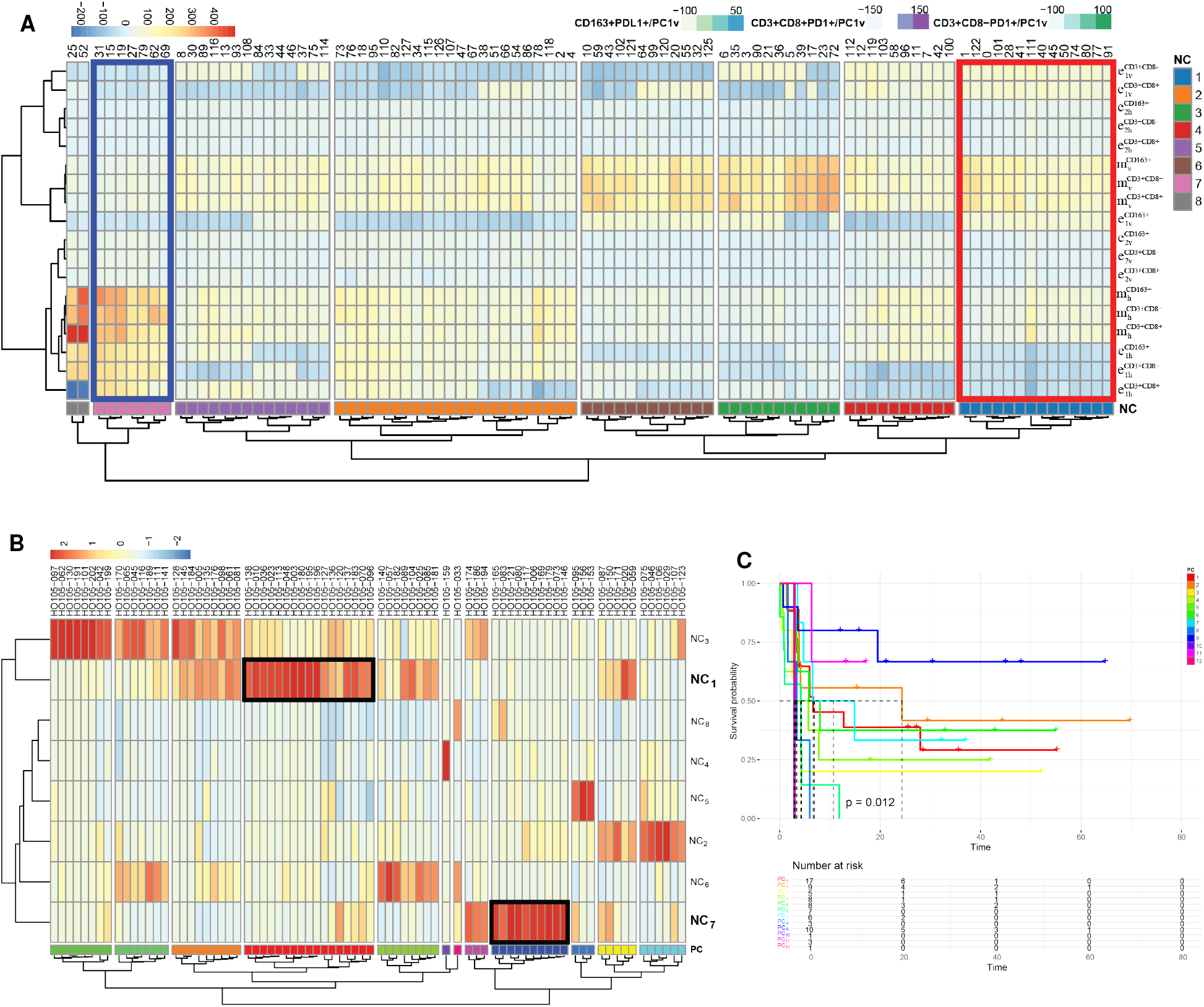
Clustering of Node Features and Patient Groups with Survival Analysis: **(A)**: Heatmap generated from the feature by node matrix, showing the average values of each feature (rows) across the node numbers (columns). Hierarchical clustering is performed using Euclidean distance with the 8 clusters, named *NC* by different colors. **(B)**: Heatmap of the *NC* frequency by patient matrix, column-scaled and clustered using Euclidean distance-based hierarchical clustering, showing the average *NC* frequency (rows) for each patient (columns). The patient clusters (*PC*s) are represented by different colors. **(C)**: Kaplan-Meier survival analysis for each *PC*.

**Figure 7:**
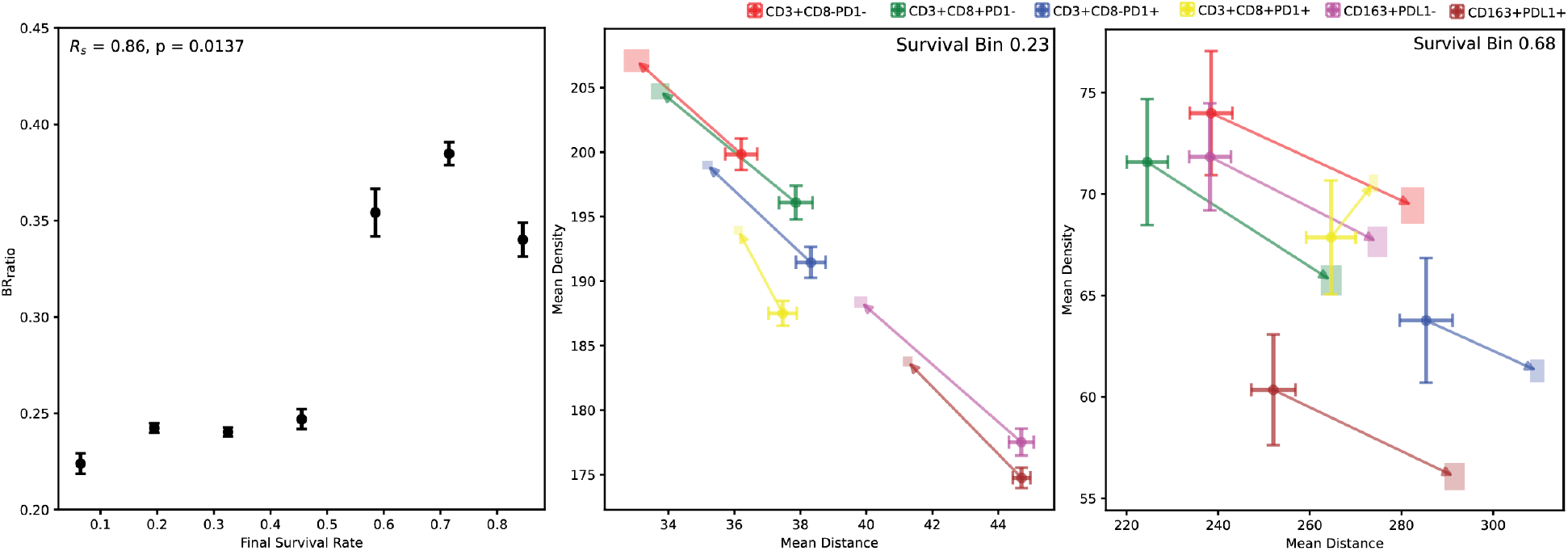
Survival Analysis Using TR and BR Images: With 628 shuffled matrices, **(Left)**: The relationship between final survival rate and the border-image ratio for corresponding patients (BR_ratio_). The border-image ratio is averaged within each bin of the final survival rate, using a bin size of 0.13. The Spearman’s Rank Correlation Coefficient (*R*_*s*_) of 0.86 indicates a strong positive monotonic relationship. **(Middle)**: Mean distance, mean density, and mean **e**_1_ for all phenotypes in the poor survival group (median survival probability = 0.23). Standard errors for distance and density are shown for each phenotype, while mean **e**_1_ is represented by arrows with error ranges (rectangles). For better visualization, we scale **e**_1_ by multiplying it by 0.05. **(Right)**: Mean distance, mean density, and mean **e**_1_ for all phenotypes in the good survival group (median survival probability = 0.68).

In the left panel of Fig. 7, a positive correlation is shown between the proportion of BR images and survival probability, indicating that *PC*s with higher survival probability have more BR images. We interpret a higher proportion of BR images as indicative of more visible border regions of the tumors. Given that the selection of TR or BR images is unbiased with respect to patients, this interesting finding implies that tumors with well-defined borders are associated with improved survival outcomes. The middle and right panels of Fig. 7 show different characteristics in the mean 𝒟𝒟 space between the poor and good survival groups. In the good survival group, all phenotypes exhibit a significantly lower mean density, indicating notably reduced concentration of malignant cells nearby. Additionally, the orientation of mean **e**_1_ on the right panel suggests that patients in the good survival group have a higher proportion of BR images, which is consistent with the result shown in the left panel. These results emphasize the importance of analyzing TR and BR images separately when studying spatial interactions between malignant and immune cells. We thus applied the two-stage HC approach using only TR (*T* = 74) and BR (*T* = 54) images, respectively, in (14). The optimal number of clusters are obtained as (*K*^∗^, *L*^∗^) = (7, 13) for TR images and (*K*^∗^, *L*^∗^) = (6, 8) for BR images.

In Fig. 8, focusing on TR images, we find that all phenotypes in the poor survival group have a higher mean density of malignant cells compared to the good survival group. This suggests that a lower mean density of malignant cells near immune cells is generally beneficial for survival. In the good survival group, active T cells (CD3+CD8+PD1- and CD3+CD8+PD1+) are positioned relatively further forward within areas of strong infiltration in the mean 𝒟𝒟 space compared to other phenotypes. However, this spatial pattern is not observed in the poor survival group. Furthermore, in the good survival group, the magnitude of mean **e**_1_ for PD1-naive T cells (CD3+CD8-PD1-) is much larger than in the poor survival group. This indicates that PD1-naive T cells exhibit a dispersed distribution in the (original) 𝒟𝒟 space, possibly reflecting their widespread presence throughout the malignant-enriched regions. Meanwhile, macrophages (CD163+PDL1- and CD163+PDL1+) are located closer to T cells in the mean 𝒟𝒟 space in the good survival group.

**Figure 8:**
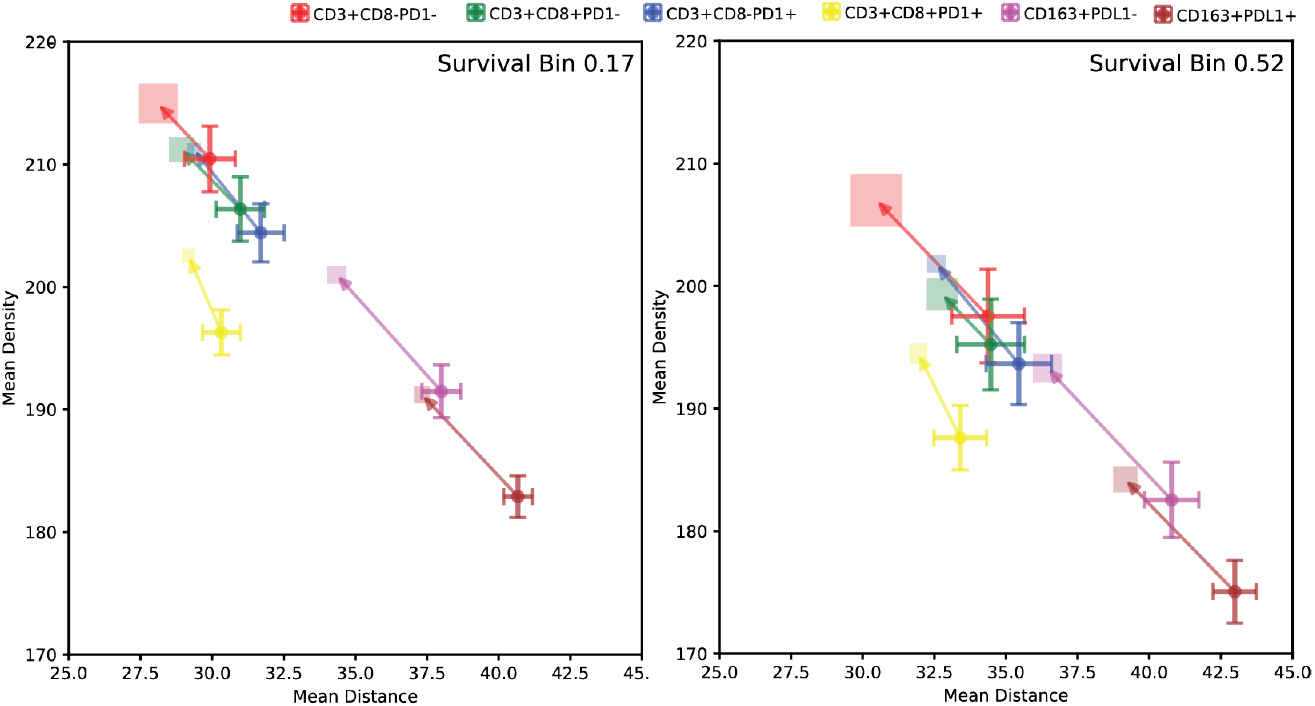
Mean Distance, Mean Density, and mean e_1_ for All Phenotypes Using TR Images: With 184 shuffled matrices, **(Left)**: Median survival probability = 0.17. **(Right)**: 0.52. For better visualization, we scale the mean **e**_1_ by multiplying it by 0.05.

In Fig. 9, we observe that the poor survival group for BR images exhibits patterns in the mean 𝒟𝒟 space similar to those seen for TR images (as shown in Fig. 8), particularly regarding the range of mean distance, mean density, and orientation of the mean **e**_1_. Moving from poor to good survival groups, immune cells are positioned further from malignant cells, implying that a high concentration of malignant cells in the border regions is particularly undesirable for survival.

**Figure 9:**
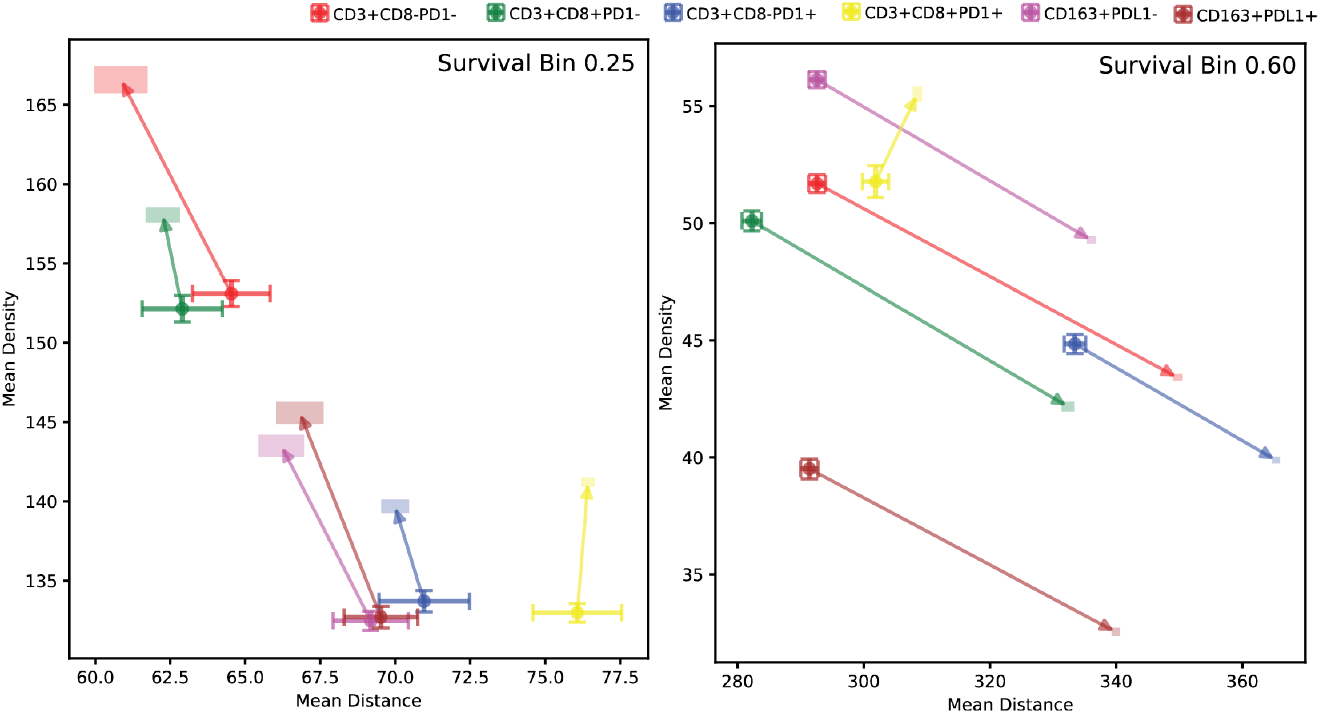
Mean Distance, Mean Density, and mean e_1_ for All Phenotypes Using BR Images: With 784 shuffled matrices, **(Left)**: Median survival probability = 0.25. **(Right)**: 0.60. For better visualization, we scale the mean **e**_1_ by multiplying it by 0.05.

## 4 Discussions

The proposed approach analyzes malignant-immune cell interactions by integrating global and local spatial scales, providing comprehensive insights into the tumor microenvironment (TME). At the global scale, immune cell infiltration is quantified using distance functions derived from the boundaries of Topological Malignant Clusters (TopMC), which are robust against noise, such as errors in cell-center extraction. Locally, malignant cell densities around individual immune cells are assessed using density functions. Principal Component Analysis (PCA) summarizes the point cloud data of each image within the distance-density (𝒟𝒟) space to characterize immune-malignant interactions. Key features, including averaged distance, averaged density, and the first two principal components, are visualized using graphs generated by Ball Mapper. These graphs reveal continuous transitions of spatial interaction patterns linked to patient survival outcomes. To ensure robustness, Ball Mapper is repeatedly applied to shuffled versions of high-dimensional feature matrices, creating multiple perspectives on the data structure. Each resulting graph of Ball Mapper is further summarized into clusters via a two-stage hierarchical clustering (HC), enabling survival analysis and uncovering associations between spatial interaction patterns and clinical outcomes.

Our analysis reveals distinct spatial patterns within the distance-density (𝒟𝒟) space between malignant-enriched (TR) and tumor-border (BR) regions. Higher proportions of BR images correlate with improved survival, suggesting that well-defined tumor borders are beneficial clinically. In TR regions, lower malignant cell density near immune cells predicts better outcomes. A notable survival-related difference involves the positioning of active T cells (CD3+CD8+PD1- and CD3+CD8+PD1+) in the mean 𝒟𝒟 space. In patients with good survival, these T cells are positioned further forward within regions of strong immune infiltration compared to other phenotypes, a feature absent in poor survival cases. Thus, optimal active T cell positioning, even when PD1 is expressed, may be crucial. Additionally, widespread distribution of PD1-naive T cells in the (original) 𝒟𝒟 space is associated with survival advantages, likely enhancing immune efficacy. Under conditions of lower nearby malignant cell density, closer proximity between macrophages and T cells in the mean 𝒟𝒟 space would correlate with better survival, potentially reflecting beneficial immune crosstalk or macrophage reprogramming within the TME [Georgoudaki et al., 2016, Tu et al., 2021].

When analyzing BR images independently, we find that patients whose border regions of the tumors exhibit similar patterns in the mean 𝒟𝒟 space with malignant-enriched regions have lower survival probabilities. This may indicate that patients whose border regions of tumors exhibit 𝒟𝒟 spatial TME patterns similar to those of malignant-enriched regions (with the presence of high malignant cell density and infiltration) are associated with poor survival outcomes [Ahn et al., 2023, 2024, Zlobec et al., 2009].

Our study demonstrates that the proposed approach robustly analyzes spatial cell distributions, consistently handling images from both TR and BR. The primary direction (**e**_1_) effectively represents a continuous spectrum from TR to BR images, providing valuable input for Ball Mapper analysis. We also reveal correlations between spatial immune phenotype patterns and patient survival. Notably, our method effectively interprets challenging immune phenotypes such as PD1+ and PDL1+ cells, previously difficult to analyze due to their low abundance in DLBCL studies [Roemer et al., 2023]. However, PCA-based analysis in the distance-density (𝒟𝒟) space remains sensitive to minor variations, especially with sparse data. Future work should focus on developing methods to reduce the impact of such variations. Overall, the robustness and adaptability of our method in interpreting diverse malignant cell distributions highlight its potential for analyzing various cancers with unique morphologies.

## Appendix

### A1 Density Function of Malignant Cells

In this section, we explain the details of how the density function *Ψ* is computed. We define a density function of the malignant cells from the reference point cloud ℛ in (1) to measure where they are densely or sparsely accumulated. Defining a square *W*_*ω*_(**r**) whose side is *ω* = 4_∅_ and center is **r** ∈ ℛ, a Gaussian function on the square is used:

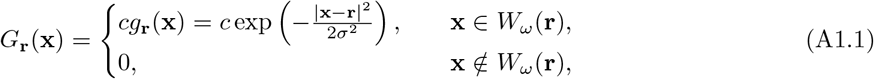

where *σ* = 0.2*ω* and *c*^−1^ is a constant 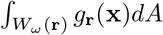, which is chosen to normalize the function over the square *W*_*ω*_(**r**). This normalization ensures that the total weight of the Gaussian function over the square equals 1, making *G*_**r**_(**x**) a probability density function within *W*_*ω*_(**r**). By scaling the function, each malignant cell in the reference point cloud ℛ contributes equally to the overall density function, independent of the size of its surrounding region. Now, the density function is defined by

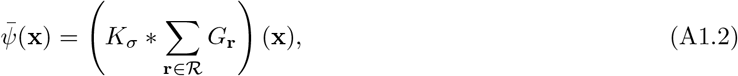

where *K*_*σ*_ is the Gaussian kernel with the standard deviation *σ* and the notation ∗ denotes the convolution operation. The value 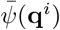 represents the density of malignant cells in the position of the immune cell **q**^*i*^ ∈ 𝒬^*i*^ in in (2). This density function quantifies how densely the malignant cells are distributed around the immune cell positions, providing insights into the local concentration of malignant cells around immune cells in the tumor microenvironment.

In order to combine *φ*(**q**^*i*^) (signed distance) and 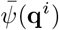 (density) in the same space, we rescale the density to the range of signed distance. This rescaling ensures that both features contribute equally when visualizing or analyzing them in the distance-density space. To do this, we first determine the global maximum (*φ*^M^) and minimum (*φ*^m^) of *φ*(**q**^*i*^) and the global maximum 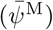 of 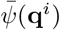 over all **q**^*i*^∈ 𝒬^*i*^ and *i* ∈ ℐacross all mIF images. Then, we normalize the density function:

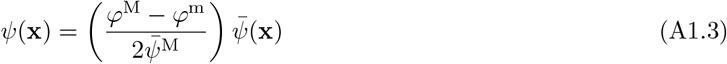

The normalization prevents one feature from dominating the analysis due to its larger numerical range and allows for an unbiased exploration of the joint effects of distance and density on immune cell behavior within the TME. From all mIF images used in this study, the values *φ*^M^ = 2079.06, *φ*^m^ = − 265.13, and 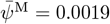 are obtained.

### A2 Algorithm of Optimal Cluster Identification

The Algorithm 1 shows a simplified step-by-step explanation of two values *P* and *S* used in optimal cluster identification.

#### Algorithm 1

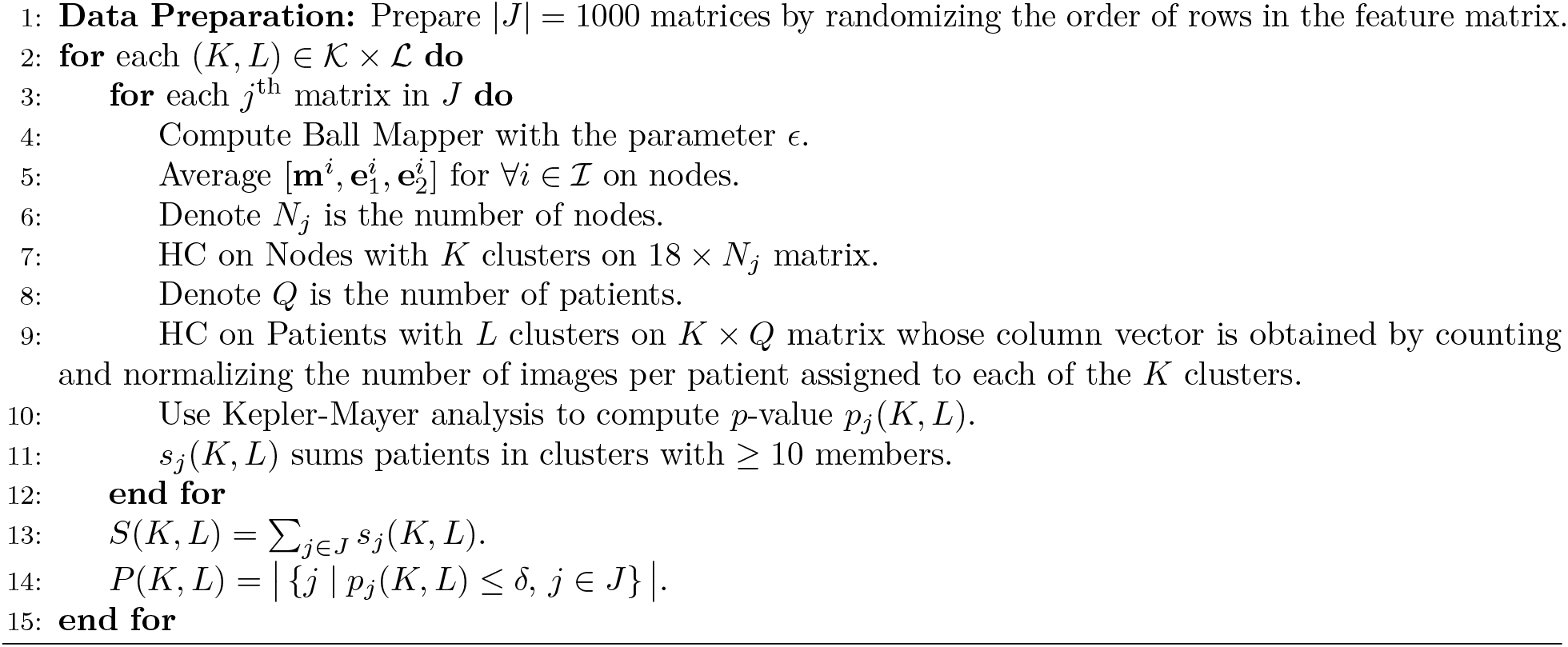

